# Plant cells at the organ surface use mechanical cues to activate a specific growth control programme

**DOI:** 10.1101/2025.08.25.672229

**Authors:** Zoe Nemec-Venza, Annamaria Kiss, Nathan German, Simone Bovio, Nadine Field, Marjolaine Martin, Claire Lionnet, Tereza Vavrdova, Qiangnan Feng, Freya de Winter, Roman Hudeček, Verena Kriechbaumer, Moritz Nowack, Charlotte Kirchhelle

## Abstract

During morphogenesis of multicellular organs, cells acquire distinct identities that meet specific functional requirements. Epidermal identity is widely considered essential for plant morphogenesis due to the role of the epidermis in both restricting and promoting growth. In the root, epidermal cells are partially covered by a protective root cap, and partially positioned at the organ surface. Here, we propose that epidermal cells at the organ surface have unique requirements for growth control due to high mechanical tension, while covered epidermal cells are mechanically shielded by the root cap. We present *in silico* and *in vivo* evidence that plants use surface mechanical cues to activate a cell-type specific growth control programme involving the small GTPase RAB-A5c, allowing them to maintain directional growth at the organ surface. Positional mechanical cues may thus be used to control expression of a sub-population of epidermal genes, linking gene regulation to surface-specific functional requirements.

## Introduction

In multicellular organisms, morphogenesis is intertwined with the establishment of different cell identities to allow the elaboration of cell types with different functions. In the context of plant development, the epidermis has received special attention thanks to its essential role in morphogenesis^1,2^. Mutants in which epidermal identity is not established properly are embryo-lethal^3,4^, and there is compelling evidence for the epidermal growth theory^1^ (i.e., the hypothesis that the epidermis determines growth of the entire organ): brassinosteroid and ethylene signalling within the epidermis is sufficient to restrict growth within the shoot^2,5^. However, in the root the epidermis can both promote growth (in longitudinal direction) and restrict growth (in radial direction)^6,7^.

Growth control necessarily involves the controlled modification of cell wall mechanical properties. In aerial organs like shoots, leaves, and the shoot apical meristem, the epidermis is generally considered to be under tension and growth-limiting^2,8–10^, and plant tissues have consequently been modelled as shells, with the outer epidermal wall being the load-bearing structure^8,11^. In line with such models, the outer epidermal cell wall is thicker and/ or stiffer than internal walls in many plant organs^1^, including the root^12^. Hormones such as brassinosteroids modulate growth through modifying cell wall properties within the epidermis^13–15^. Effects on directional growth such as those observed in the root can be explained through a direct effect on microtubule organization^16–18^, which in turn determine the organization of cellulose in the cell wall to promote directional growth^19,20^. In addition to cellulose orientation, plants also employ an additional growth control pathway mediated by the small GTPase RAB-A5c, which defines a transport pathway to cell edges and is expressed in epidermal cells^21,22^. Epidermal growth control can thus involve tissue-specific regulation of generic growth control pathways as well as cell-type specific expression of specialized growth control pathways. It has been speculated that the acquisition of epidermal identity may be linked to its mechanical status^23^, suggesting a reciprocal connection between cell identity and cell mechanics within the epidermis.

Here, we use roots to explore the relationship between gene expression, cell mechanics, and growth control within epidermal cells. In root tips, the epidermis is partially covered by the root cap, a protective cell layer which undergoes programmed cell death (PCD^24^), allowing the comparison between covered and uncovered epidermal cells. We show that RAB-A5c is expressed primarily in epidermal cells at the surface of developing roots, while it is inhibited by root cap cover. We present *in silico* and *in vivo* data that causally link mechanical stress to the expression of RAB-A5c. We propose that RAB-A5c expression is triggered in cells at the organ surface through high mechanical stress, where it is needed to regulate cell growth in this unique mechanical niche.

## Results

### RAB-A5c is expressed in growing cells at the root surface

We previously reported that in lateral roots, *pRAB-A5c::YFP:RAB-A5c* (YFP:RAB-A5c) was primarily expressed in meristematic epidermal cells, and expression decreased with differentiation^21^. On lateral root transverse and longitudinal cross-sections, it was apparent that even within the meristem, YFP:RAB-A5c was not uniformly expressed, but was largely absent in epidermal cells covered by the root cap (Figure 1A-C,E). The root cap ends in different positions for each cell file^25^ (Figure 1A,B,I,J), but 2D and 3D quantitative analysis of YFP:RAB-A5c expression along epidermal cell files using the root cap as a landmark revealed a consistent pattern: YFP:RAB-A5c expression was low in covered epidermal cells with a mild increase towards the end of the root cap, but sharply increased at the root cap boundary (Figure 1B,G, S1A). This expression pattern was drastically different from that of the related Rab-GTPase *pRAB-A2a::YFP:RAB-A2a*^26^, which was uniformly expressed along epidermal cell files, irrespective of root cap cover (Figure 1D,F,H, S1B,C). In contrast to *pRAB-A5c::YFP:RAB-A5c*, YFP:RABA5c expressed under the ubiquitous promoter pUBIQUITIN10 (*pUB10::YFP:RAB-A5c*) did not show surface-specific fluorescence (Figure S1D), indicating that the YFP:RAB-A5c surface pattern was transcriptionally controlled.

**Figure 1:**
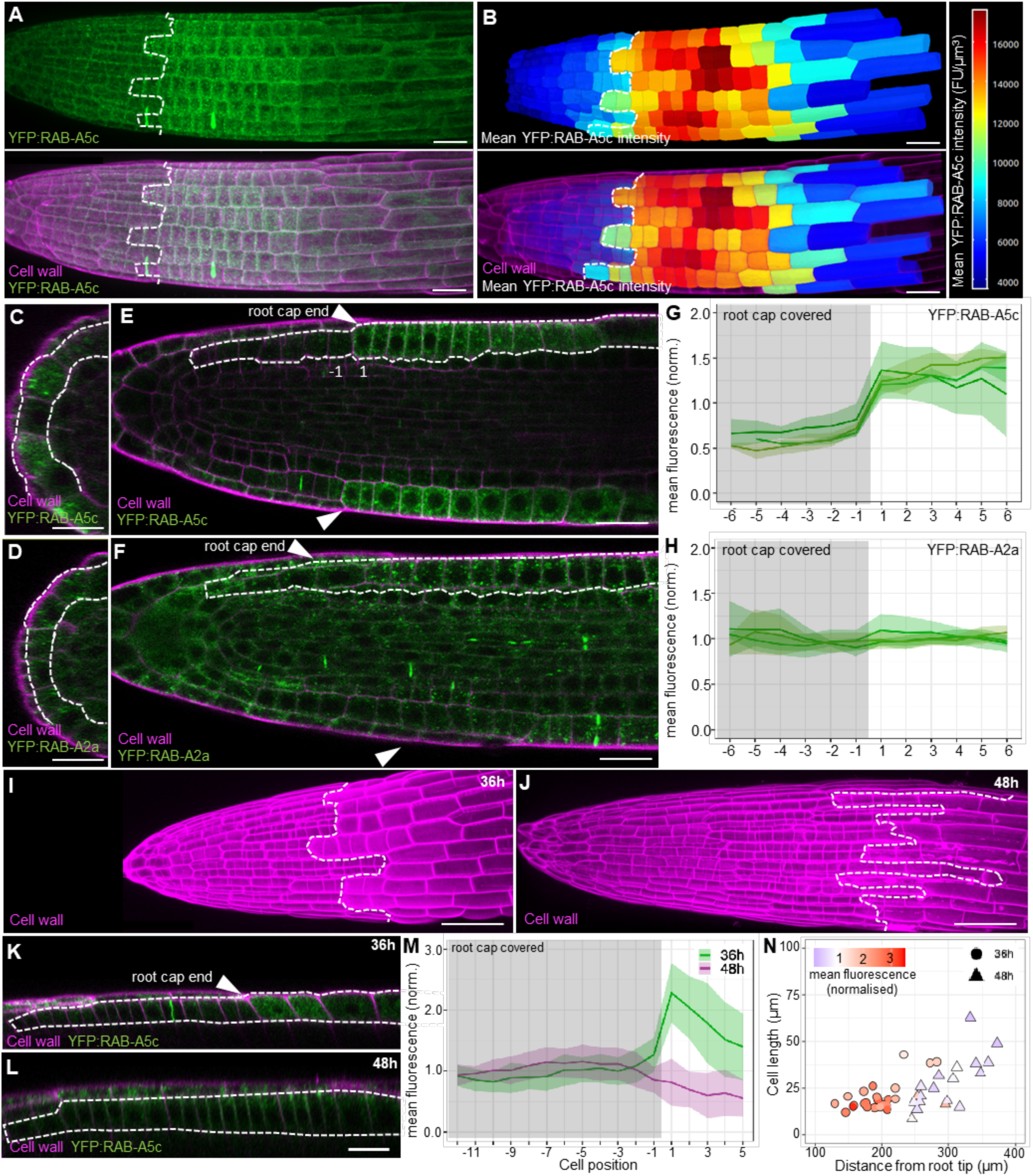
RAB-A5c expression is limited to epidermal cells at the organ surface. **(A)** Maximum intensity projection of a confocal laser scanning microscopy (CLSM) stack of a lateral root expressing YFP:RAB-A5c stained with the cell wall marker propidium iodide (PI). Dashed line indicates the end of the root cap. **(B)** MorphoGraphX 3D segmentation of epidermal cells in the lateral root shown in (A). Heat map depicts mean fluorescence intensity of YFP:RAB-A5c. Dashed line indicates the end of the root cap. FU = Fluorescence Units **(C-F)** YZ cross-sections (C,D) or XY midplane optical sections (E,F) of CLSM stacks from PI-stained lateral roots expressing either YFP:RAB-A5c (C, E, from root shown in A) or YFP:RAB-A2a (D,F, Fig S1). Arrow heads indicate the end of the root cap, dashed line surrounds epidermal cells. Numbers indicate position of the first covered (-1) and the first uncovered (1) epidermal cells. **(G,H)** Mean fluorescence intensity of YFP:RAB-A5c (G) or YFP:RAB-A2a (H) per cell along epidermal cell files from roots such those shown in (A-F). Fluorescence was quantified in 3D along 3-5 individual cell files per root, cell files were aligned based on the position of the root cap, with the last covered cell labelled as -1 and the first uncovered cell labelled as 1. Ribbon plots represent average fluorescence +/- 1SD for each root. N = 3 roots. **(I,J)** Maximum intensity projections of PI-stained primary roots aged 36h (I) or 48h (J). Dashed line indicates the end of the root cap. **(K,L)** XY optical section through 36h (K) and 48h (L) old PI-stained primary roots expressing YFP:RAB-A5c. Arrow heads indicate the end of the root cap, dashed line surrounds epidermal cells. Note the meristem is fully covered by the root cap at 48h. **(M)** Mean fluorescence intensity of YFP:RAB-A5c per cell along epidermal cell files in 36h and 48h old primary roots such those shown in (I-L). N=23 and 21 cell files from 5 roots (36h and 48h, respectively). Cell files were aligned based on root cap position, with the last covered cell labelled as -1 and the first uncovered cell labelled as 1. Ribbon plots represent average fluorescence +/- 1SD. **(N)** Plot showing the mean fluorescence intensity, cell length, and distance from the root tip of the first uncovered epidermal cell in 36h and 48h old primary roots such as those shown in (I-L). Scale bars: 20µm.

In primary roots the root cap progressively covers the entire meristem within the first two days after germination (Figure 1I,J). In primary roots with a partially uncovered meristem (36h old), we observed a strong peak of YFP:RAB-A5c expression in the first uncovered cells (Figure 1K,M), matching observations in lateral roots (Figure 1C,E). In primary roots with an almost completely covered meristem (48h old), YFP:RAB-A5c expression did not increase at the end of the root cap, although some expression was observed within covered meristematic cells (Figure 1L,M). This suggests YFP:RAB-A5c expression is only activated in uncovered cells when these are meristematic. In line with this, first uncovered cells at 48h were on average both further away from the root tip and longer than at 36h (Figure 1N), suggesting they had moved out of the meristematic zone. Notably, first uncovered cells in 36h old roots also showed less activation with increasing length and distance from the root tip, indicating a “window of competence” for *YFP:RAB-A5c* activation within the meristem, irrespective of root age (Figure 1N).

### Root cap removal induces YFP:RAB-A5c expression

To test whether *YFP:RAB-A5c* activation was functionally linked to root cap death, we expressed NAC46, a transcription factor which is sufficient to initiate programmed cell death^27^, under the control of a root-cap specific, dexamethasone (Dex)-inducible expression system (*pSMB:GRLhG4>>pOP6-ANAC046-P2A-mtagBFP2-NLS;* short *pSMB:GRLhG4>>NAC46-BFP*). After 48h treatment with Dex (inducing condition), we observed a drastic reduction of root cap length in comparison to DMSO-treated controls, with none or few root cap cells covering the meristematic epidermis (Figure 2A,B,I,J). Despite the resulting shift of the first uncovered cell towards the root tip in Dex-treated roots, YFP:RAB-A5c was expressed in comparable levels in the first uncovered cell of DMSO and Dex-treated roots (Figure 2C,D). In contrast to DMSO-treated controls, expression levels in uncovered cells of Dex-treated roots increased further with increasing distance from the root cap, (Figure 2D), resulting in overall higher YFP:RAB- A5c expression levels (Figure S2D).

**Figure 2:**
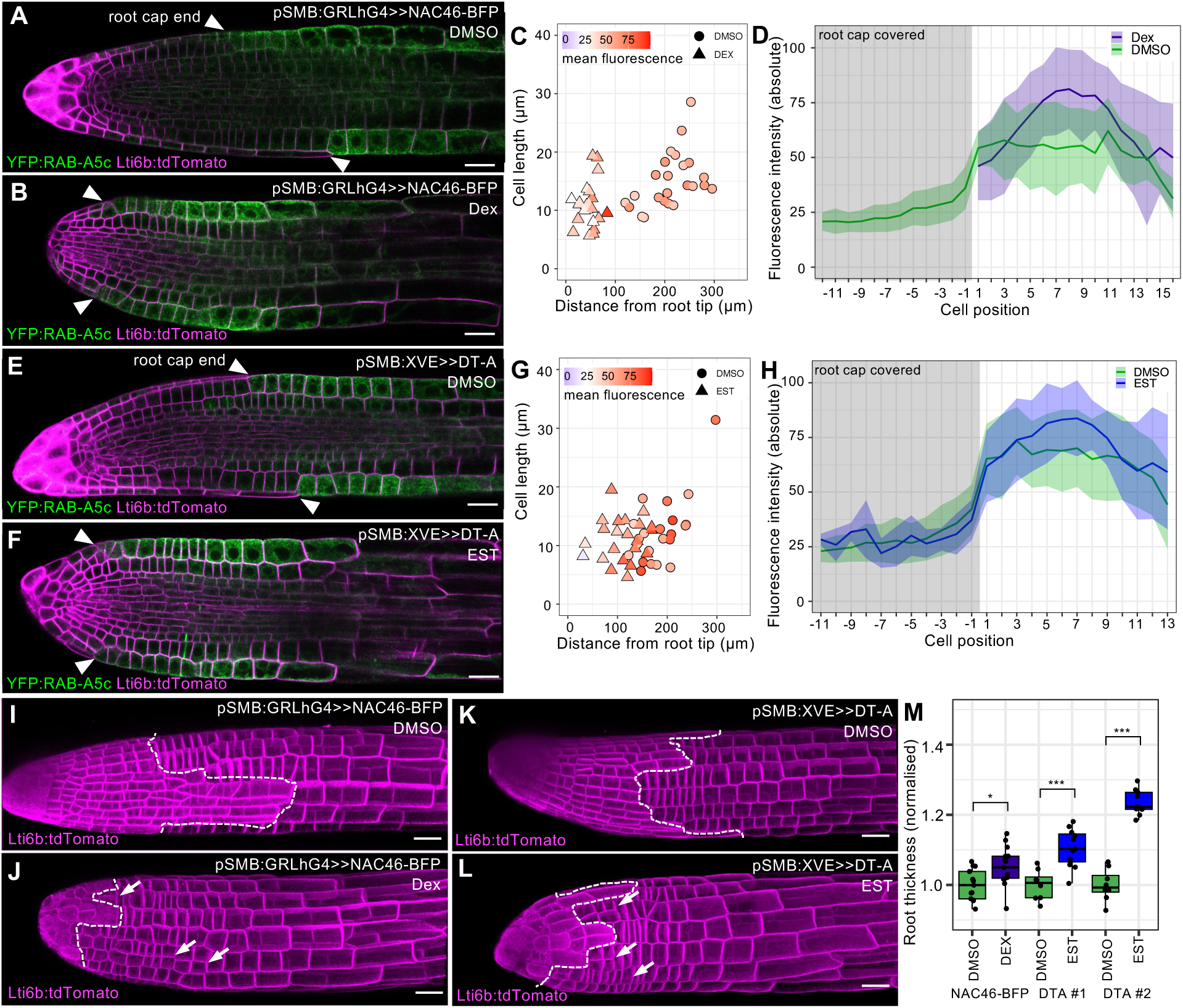
Removal of root cap cells induces YFP:RAB-A5c expression. (A,B) XY optical sections of CLSM stacks from lateral roots co-expressing YFP:RAB-A5c, Lti6b:tdTomato, and Dexamethasone (Dex)- inducible *pSMB:GRLhG4>>NAC46-BFP* after 48h treatment with DMSO as a control (A) or 1µM Dex (B) to induce root cap death. **(C)** Plot showing the mean fluorescence intensity, cell length, and distance from the root tip of the first uncovered epidermal cell in lateral roots such as those shown in (A,B). **(D)** Mean fluorescence intensity of YFP:RAB-A5c per cell along epidermal cell files lateral roots such those shown in (A,B). N=23 (Dex) and 24 (DMSO) cell files from 5 roots. Fluorescence was quantified in 2D, cell files were aligned based on the position of the root cap, with the last covered cell labelled as -1 and the first uncovered cell labelled as 1. Ribbon plots represent average fluorescence +/- 1SD. (**E,F)** XY optical sections of CLSM stacks from lateral roots co-expressing YFP:RAB-A5c, Lti6b:tdTomato, and Estradiol (Est)- inducible *pSMB:XVE>>DT-A* #1 after 48h treatment with DMSO as a control (E) or 5µM Est (F) to induce lateral root cap death. **(G)** Plot showing the mean fluorescence intensity, cell length, and distance from the root tip of the first uncovered epidermal cell in lateral roots such as those shown in (E,F). **(H)** Mean fluorescence intensity of YFP:RAB-A5c per cell along epidermal cell files lateral roots such those shown in (E,F). N=22 cell files from 5 roots for both Est and DMSO. Fluorescence was quantified in 2D, cell files were aligned based on the position of the root cap, with the last covered cell labelled as -1 and the first uncovered cell labelled as 1. Ribbon plots represent average fluorescence +/- 1SD. **(I-L)** MorphoGraphX renderings of lateral roots such as those shown in (A,B,E,F). Dashed line indicates the end of the root cap, arrows indicate misplaced longitudinal divisions. **(M)** Box plot of lateral root thickness from roots such as those shown in (I,L). Root thickness was normalized against DMSO controls for each genotype. N = 8 (DT- A#1 DMSO, DT-A#2 DMSO), 9 (DT-A#2 Est), 11 (DT-A#1 Est), 13 (NAC46 DMSO), 15 (NAC46 DEX). Significant differences in thickness are indicated as follows: * p<0.05; *** p<0.001, Two-way ANOVA and post-hoc Tukey test. Scale bars: 20µm.

To understand whether *YFP:RAB-A5c* activation in the epidermis was linked to signals associated with root cap PCD, we quantified *YFP:RAB-A5c* expression in the *sombrero (smb)* mutant background, in which root cap PCD is delayed^28^. We still detected YFP:RAB-A5c in uncovered epidermal cells of 36h primary roots of *smb* plants (Figure S2A-C), indicating that PCD was not required to trigger YFP:RAB-A5c expression. To test this hypothesis further, we expressed a diphtheria toxin A-chain (DT-A) gene^29^ under the control of a root cap-specific, estradiol (EST)-inducible expression system (*pSMB:XVE>>DT-A*) as a PCD-independent strategy to induce root cap death. After 48h treatment with either EST (inducing condition), we observed a strong reduction in root cap length in comparison to the DMSO control (Figure 2E,F,K,L, S2F,G). *YFP:RAB-A5c* expression in covered cells and the first uncovered cell in EST-treated and control plants was comparable, followed by an increase of peak fluorescence in EST-treated plants only, similar to the pattern observed after induced PCD (Figure 2G,H, S2E).

Taken together, these results indicated that unlike for processes like hormone patterning in the root^30^, it was not PCD-related signalling, but rather the removal of the root cap *per se* that triggered *YFP:RAB-A5c* expression. Notably, root cap removal resulted in geometric changes within the root meristem, with a significant increase of root thickness of 5.6% in *pSMB:GRLh4>>NAC46-BFP* and 11.5% or 25.9% in *pSMB:EST>>DT-A* roots, respectively (Figure 2M). The increased root diameter coincided with changes in cell division pattern: we observed anticlinal radial divisions in the epidermis, which are normally absent (Figure 2I-L, S2F-G). In other organs, tissue tension can pattern cell division plane orientation^31^. We therefore hypothesized that maximum tension is born by the root cap in wild-type roots, but transferred to the epidermis when the root cap is removed either through PCD or toxin expression.

### Root surface cells bear maximum tension *in silico*

To explore root tension patterns, we built 2D finite element models of lateral root cross-sections (Figure 3). To parameterize the model, we measured cell wall thicknesses in serial block face scanning electron micrographs of a wild-type lateral root (Figure 3A-C, S3A-C). Similar to primary roots^12^, we found that the cell wall at the organ surface was approximately twice as thick as inner cell faces shared between two cells, both in epidermal and root cap cells (Figure 3D,E). We quantified cell sizes from confocal microscopy images of wild-type lateral roots within the zone where both covered and uncovered cells co-exist, and *YFP:RAB-A5c* is activated (Figure S3D-G). We constructed idealized lateral root cross-sections representing three cases: (a) a lateral root covered by a root cap, (b) a lateral root from which the root cap was removed without increasing surface wall thickness (e.g. corresponding to DT-A toxin expression), and (c) a lateral root with uncovered epidermis cells in which the outer cell wall was thickened, in line with our physiological data (Figure 3F). We treated cell walls as a linear elastic material with a Young’s modulus (E) and Poisson ratio of 0.3 and inflated cells with a turgor pressure of P=0.005E, so that the system was kept in a small deformation regime. In simulations with uniform cell wall properties, maximum tension was consistently highest in outermost tissue layer for all three cases (Figure 3G), similar to the elongation zone of primary roots^12^. We also examined the relative change in cell area before and after applying turgor pressure, and observed that covered epidermal cells (case a) or uncovered ones (case c) were similarly deformed, whereas the removal of the root cap without wall thickening (case b) caused a drastic increase in cell area (Figure 3H, S3H-J). When the root cap was present, it was exposed to maximum tension, but did not undergo maximum area change (Figure 3H, S3H-J). We confirmed this prediction experimentally by releasing turgor pressure in lateral roots through osmotic treatments, which reveals pressure-driven area changes in the form of shrinkage^32^. Both volumetric and cross-sectional area shrinkage varied substantially from root to root, but we consistently observed that root cap cells shrank less than covered and uncovered cells, as predicted in our model (Figure 3I-J, S3K-R).

**Figure 3:**
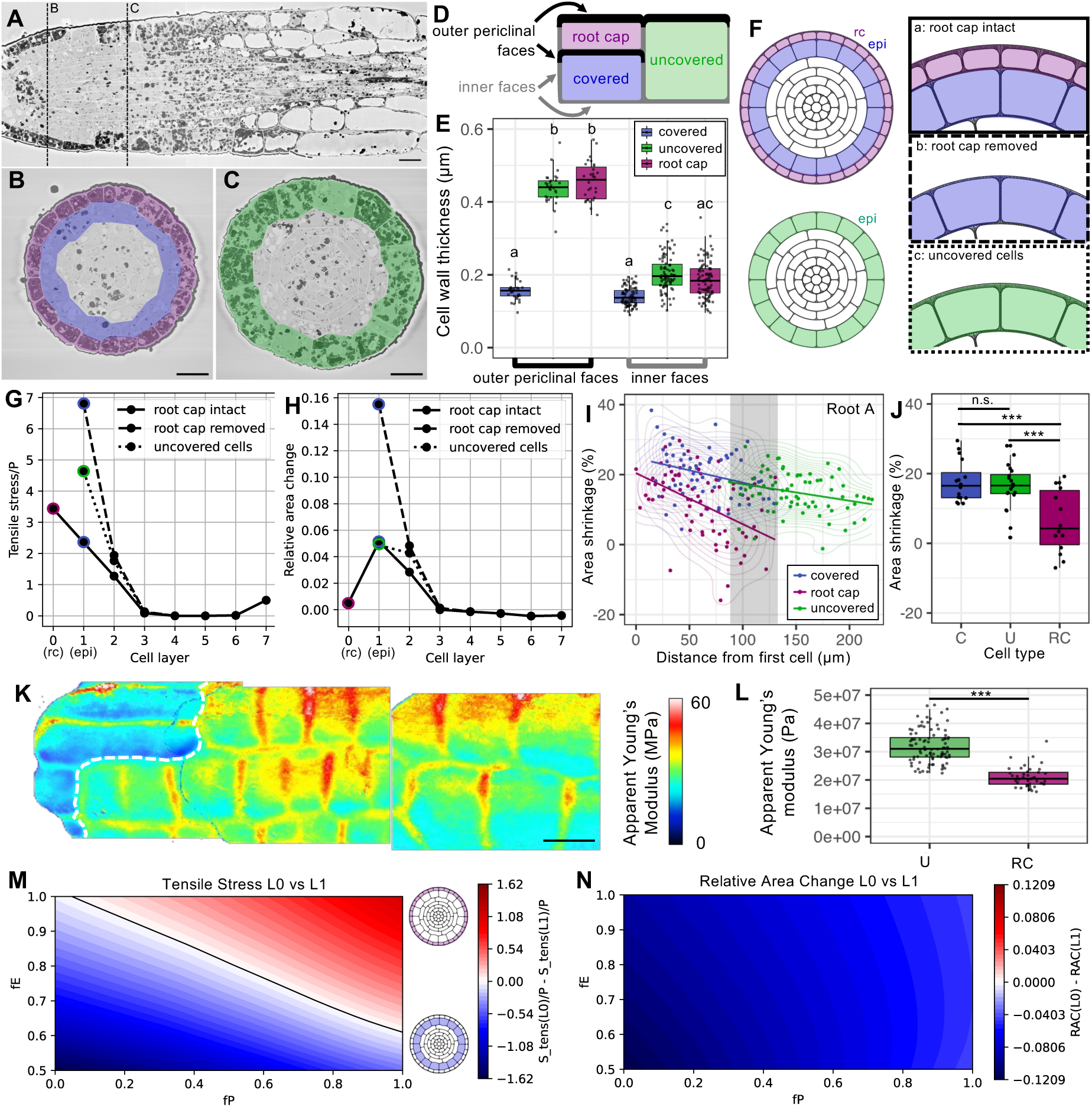
The organ surface bears maximum tension in lateral roots. (A-C) Longitudinal (A) and transverse (B,C) cross-sections of a serial block-face scanning electron micrograph (SBF-SEM) of a PFA-fixed wild- type lateral root. Root cap cells are false-coloured purple, covered epidermal cells blue, uncovered epidermal cells green. **(D)** schematic depiction of cell faces in root cap and epidermal cells such as those shown in (B,C). **(E)** Cell wall thickness at different cell faces as shown in (D), quantified from SBF-SEM sections such as those shown in (B,C). Same letters indicate no significant difference (p≥0.05), different letters a significant difference (p<0.05), Two-way ANOVA and post-hoc Tukey test. **(F)** Reference configuration of computational models of lateral root cross-sections. **(G,H)** Tensile stress (G) and relative cross-sectional area change (H) in all cell layers of the model cases a,b, and c shown in (F), after inflation with P/E=0.005. **(I)** Cross-sectional area shrinkage after plasmolysis in root cap, covered epidermal, and uncovered epidermal cells as a function of distance from the root tip. Area between the first uncovered and last covered cell is shaded in grey. **(J)** Box plot of cross-sectional area shrinkage of cells in the shaded area shown in (I). Significant differences are indicated as follows: n.s. p≥0.05; *** p<0.001, One-way ANOVA and post-hoc Tukey test. **(K)** Atomic Force Microscopy (AFM) map of a lateral root in water. Dashed line indicates the end of the root cap. **(L)** Apparent Young’s modulus of cell edges in uncovered epidermal (U) and root cap (RC) cells from scans such as (K). N = 41 (RC), 94 (U) from 4 roots. **(M,N)** Map of the difference in tensile stress (M) and relative cross-sectional area change (N) between the root cap (L0) and covered epidermis (L1) in response to changes in turgor pressure (fP) and elastic modulus (fE) within the L0 layer. Blue color indicates that the respective value is higher in the epidermis, red color indicates higher values in the root cap. Scale bars = 10µm

Models with uniform cell wall properties and turgor were thus sufficient to explain experimental observations. However, in atomic force microscopy measurements, we observed that the root cap’s stiffness (Apparent Young’s modulus) was 34.2% lower than the meristem (Figure 3K,L). This effect may either be due to differences in cell wall elasticity or in turgor pressure, which can influence the apparent Young’s modulus and is likely affected by vacuole collapse and plasma membrane permeabilization in root cap cells preceding programmed cell death^33^. We changed turgor pressure and/or Young’s modulus within the root cap (L0) layer *in silico* (case a), and found that reduction of the Young’s modulus by at least 40% shifted the maximum tension to the L1 layer, as did a combination of turgor pressure and Young’s modulus reduction (Figure 3M, black line, S4A,B). Such changes did not substantially alter relative area change within the different tissue layers, even when maximum tension was shifted from the L0 to the L1 layer (Figure 3N, S4C,D). Taken together, these results suggest that cells at the root organ surface experience maximum tensile forces, necessitating mechanical reinforcement. Furthermore, the root cap can shield covered meristematic cells from maximum tension, but changes in root cap turgor pressure and/or mechanical stiffness can shift the tension maximum to the epidermal layer or maintain the two layers at a similar tension level.

### Global variation in cell wall tension modifies YFP:RAB-A5c expression

Mechanical signals have been linked to gene expression control in the shoot apical meristem^34^ and leaf^35^. We therefore tested whether the mechanical status of surface cells was linked to activation of *YFP:RAB-A5c* in roots. Uniform reduction (or increase) of turgor pressure across all cell layers linearly reduced (or increased) tensile stress in the L0 and L1 layers in the model (Figure 4A,B). To emulate the same effect experimentally, we treated YFP:RAB-A5c-expressing plants grown on solid ½ MS medium for six hours with iso-osmotic (½MS), hyper-osmotic (½MS+200mM sorbitol), or hypo-osmotic (water) solutions (Figure 4C-E). Iso-osmotic treatment globally preserved the YFP:RAB-A5c pattern of untreated plants (Figure 4D,F, S5A,B). Hyper-osmotic treatment, which is expected to reduce turgor pressure and cell wall tension, reduced YFP:RAB-A5c levels in all epidermal cells compared to control conditions (Figure 4C-D,F-G). Furthermore, the gradient in covered epidermal cells was reduced (Figure S5C,D), causing a significantly steeper activation at the root cap boundary (Figure 4F,H). Hypo-osmotic treatment had the inverse effect, mildly increasing YFP:RAB-A5c in covered and uncovered cells (Figure 4D-G) and causing a steeper gradient of fluorescence in covered cells (Figure S5D,E), although this effect was more subtle and did not affect the activation at the root cap boundary (Figure 4H). These findings suggest that levels of YFP:RAB-A5c in both covered and uncovered cells are sensitive to changes in turgor pressure and/or cell wall tension. Covered cells close to the root cap end were the most sensitive to changes, consistent with the notion that these were closest to each other in terms of tension. To separate cell wall tension from osmotic effects, we also grew plants on ½ MS plates with high amounts of agar, a treatment previously described to reduce cell wall tension without affecting turgor pressure^36^ (Figure S5F-I). YFP:RAB-A5c levels were significantly reduced in lateral roots grown on plates supplemented with 2.5% agar compared to 0.8% agar in both covered and uncovered cells (Figure S5H,I), indicating that cell wall tension could be directly linked to YFP:RAB-A5c expression.

**Figure 4:**
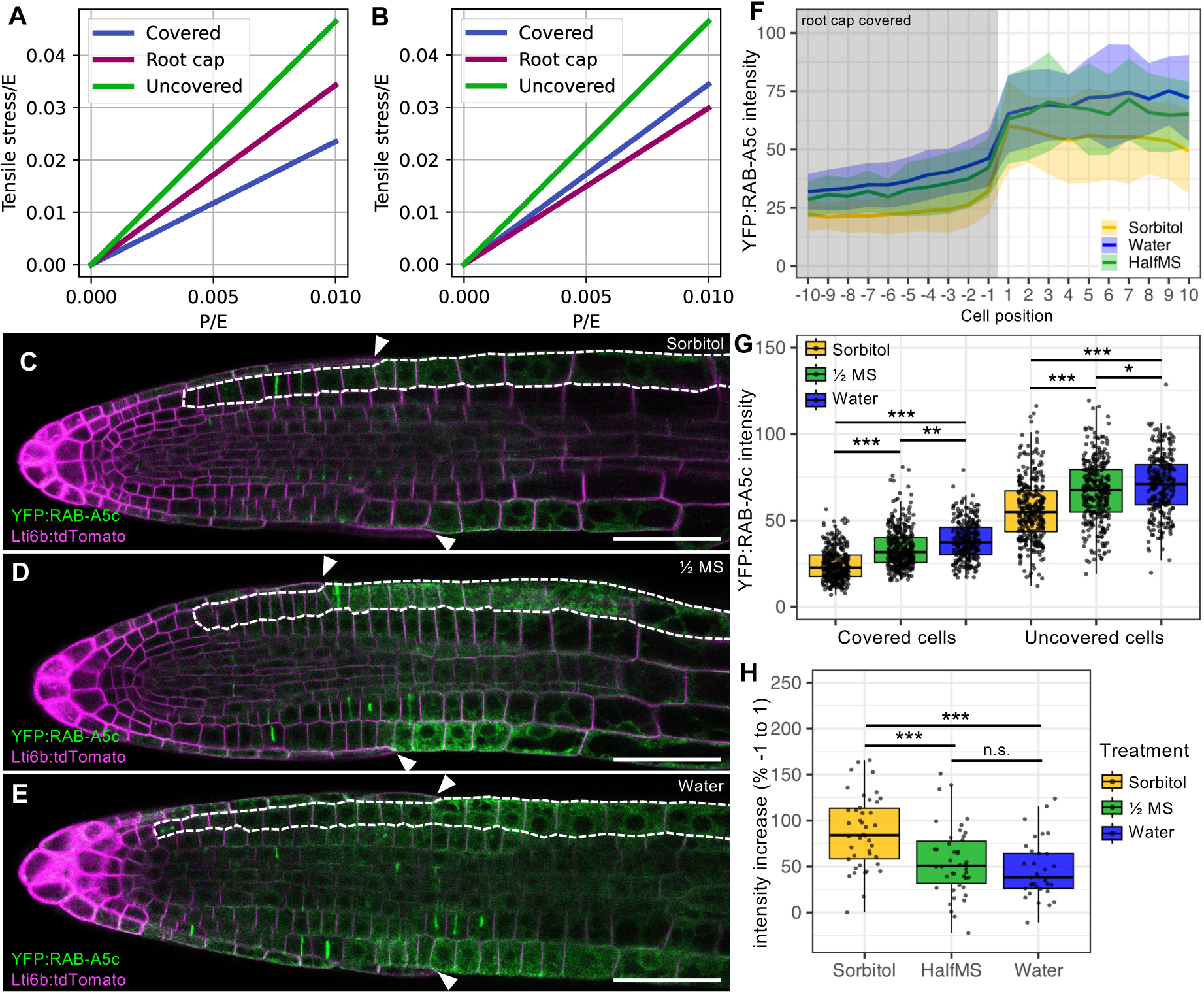
RAB-A5c expression is sensitive to global changes in tissue tension. (A,B) Tensile stress in computational models of the configuration shown in Figure 3F, case a (root cap intact) in response to changes in turgor pressure (P/E) across all tissue layers. Elastic modulus E was uniform across all tissue layers (A) or reduced by 50% in the root cap compared to the remaining tissue layers (B). **(C-E)** XY optical sections of CLSM stacks from lateral roots co-expressing YFP:RAB-A5c and Lti6b:tdTomato after 6h treatment with liquid ½MS+200mM Sorbitol (C), ½ MS (D), or water (E). Dashed line surrounds epidermal cells in one cell file. Arrowheads indicate end of the root cap. **(F)** Mean fluorescence intensity of YFP:RAB- A5c per cell along epidermal cell files lateral roots such those shown in (C-E). N=33 (Water), 43 (Sorbitol, HalfMS) cell files from 10 roots (2 replicates). Fluorescence was quantified in 2D, cell files were aligned based on the position of the root cap, with the last covered cell labelled as -1 and the first uncovered cell labelled as 1. Ribbon plots represent average fluorescence +/- 1SD. **(G,H)** Box plots of average YFP:RAB- A5c intensity in covered and uncovered cells (G) or intensity increase between cell -1 and 1 (H) from data shown in (F). Significant differences are indicated as follows: n.s p≥0.05; * p<0.05; ** p<0.01; *** p<0.001, One- or Two-way ANOVA and post-hoc Tukey test. Scale bars: 50µm.

### Local increases in cell wall tension induce YFP:RAB-A5c expression

Cell ablations have been used previously to vary cell wall tension pattern locally through increasing tension around the wound site^37^, an effect we also observed by ablating root cap cells *in silico* (Figure 5A-D). To test whether cell ablations trigger *YFP:RAB-A5c* expression *in vivo*, we ablated root cap cells by laser ablation, and examined YFP:RAB-A5c before, immediately after, 6h after, and 24h after the ablation (Figure S6A-H). We noted that epidermal cells of interest beneath the ablation were strongly photobleached up to 6h after the ablation, which hampered systematic assessment of YFP:RAB-A5c fluorescence. We nevertheless occasionally observed increased YFP:RAB-A5c levels after 6h and 24h (Figure S6C,D,G,H). To minimise photobleaching/-toxicity effects, we ablated both root cap and epidermal cells and observed fluorescence in cells adjacent to the wound site, outside of the laser line (Figure 5E-H, M-O). Despite the mild photobleaching, YFP:RAB-A5c fluorescence systematically increased after 6h and 24h in adjacent to the wound, in both covered and uncovered epidermal cells (Figure 5I-LP) as well as in root cap and cortex cells (Figure 5M-O, S6I-P). Thus, YFP:RAB-A5c expression was not strictly linked to the epidermal lineage, but could be triggered by a sudden increase in cell wall tension in other cell types, and even in cells already at the organ surface.

**Figure 5:**
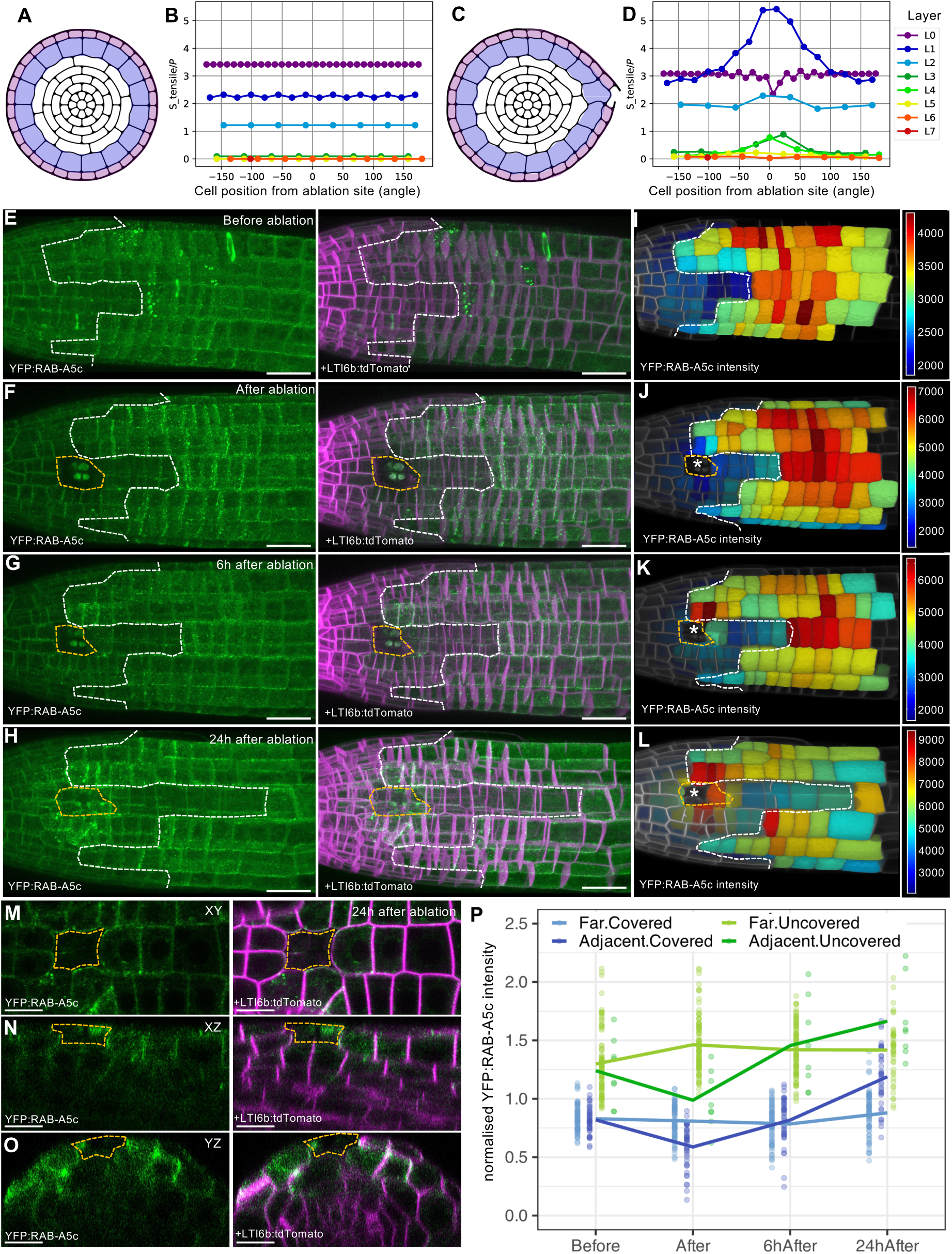
RAB-A5c expression is activated around ablation sites. (A-D) Computational models (A,C) and predicted tensile stress (B,D) in intact roots (A,B) and roots where one root cap cell was ablated (C,D). Note maximum tensile stress is transferred from the L0 to the L1 later in the vicinity of the ablated cell. **(E-L)** Maximum intensity projections of CLSM stacks (E-H) and MorphographX volumetric fluorescence intensity maps (I-L) of a lateral root co-expressing YFP:RAB-A5c and Lti6b:tdTomato before and immediately after, 6h, and 24h after ablation within the root tip. Dashed white line indicates end of the root cap, dashed yellow line and asteriscs indicates the ablation site. **(M-O)** XY, XZ, and YZ sections around the wound site of the root shown in H,L, 24h after ablation. Dashed yellow line indicates the ablation site. **(P)** YFP:RAB-A5c intensity normalized by average root fluorescence from cells in 7 roots such as that shown in (E-O). Cells were divided into four groups: covered epidermal cells adjacent to (n= 40) or far from (n=71) the wound site, and uncovered epidermal cells adjacent to (n= 8) or far from (n=94) the wound site. Scale bars: 20µm.

### RAB-A5c is required to maintain directional growth in surface cells

We previously demonstrated that RAB-A5c contributes to directional growth control independently of cellulose organisation^22^. Conditional inhibition of RAB-A5c via expression of the dominant-negative RAB-A5c[N125I] variant caused severe morphological defects at the cell- and organ scale in young lateral roots, however this phenotype got progressively less severe in older lateral roots, in which the root cap covers an increasing proportion of the meristem^21^. To test whether *RAB-A5c* activation by cell wall tension is required specifically in meristematic surface cells, we explored whether altering cell wall tension had phenotypic consequences for lateral root morphogenesis. Growing plants on plates containing ½MS+200mM sorbitol for 2d or 3d significantly reduced YFP:RAB-A5c levels in uncovered meristematic cells (Figure 6A-C, S7). We used this treatment to reduce YFP:RAB-A5c levels in plants expressing the dominant-negative RAB-A5c[N125I] protein under the control of the Dexamethasone-inducible LhGR system^21^ (*pRPS5a>>Dex>>RAB-A5c[N125I]*, hereafter *RAB-A5c[NI]*). We quantified lateral root thickness as a proxy for defects in directional growth^21,22^. While treatment with ½MS+200mM sorbitol did not affect root thickness in wild-type seedlings, *RAB-A5c[NI]* plants showed severely and significantly enhanced root swelling compared to ½MS controls (Figure 6D-L). Growth defects were most severe in uncovered cells, while morphology in the region covered by the root cap was only mildly perturbed (Figure 6D- K). We conclude that RAB-A5c homeostasis driven by cell wall tension is essential for the directional growth of epidermal cells at the organ surface.

**Figure 6:**
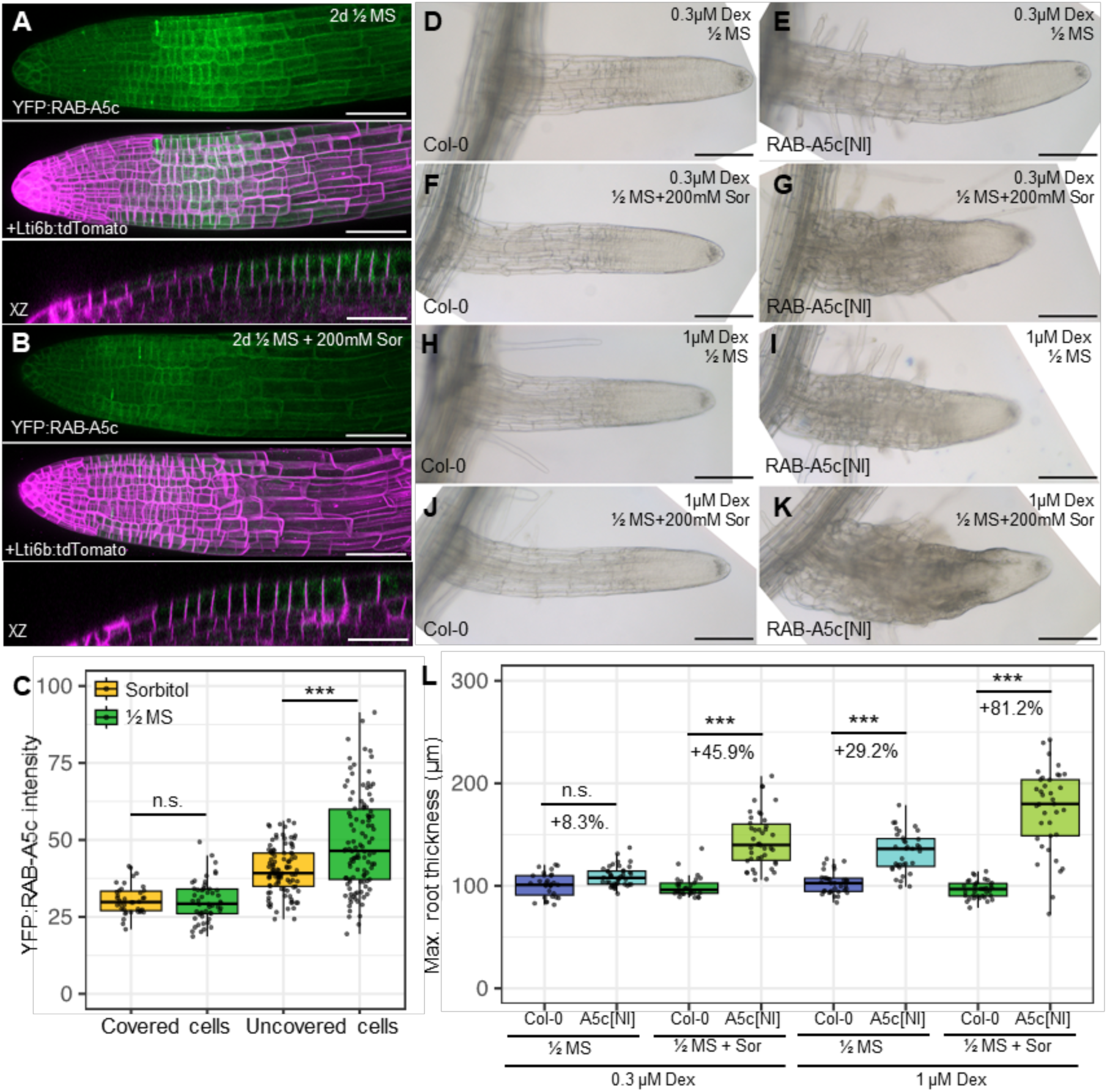
Lateral roots are sensitive to sorbitol treatment when RAB-A5c function is perturbed. (A,B) Maximum intensity projections and XZ sections of CLSM stacks of lateral roots co-expressing YFP:RAB- A5c and Lti6b:tdTomato after 2d on plates containing ½ MS (A) or 1/2MS + 200mM Sorbitol (B). **(C)** Box plot showing YFP:RAB-A5c intensity in covered and uncovered cells from roots such as those shown in (A,B). **(D-K)** Photographs of lateral roots from wild type (D,F,H,J) or RAB-A5c[NI] (E,G,I,K) plants grown for 8d on ½ MS and 3d on ½ MS (D-E, H-I) or ½ MS with 200mM Sorbitol (F-G, J-K) supplemented with\ 0.3 (D-G) or 1µM Dex (H-K). **(L)** Box plots of lateral root thickness of roots such as those shown in (D-K). n= 33-37 roots per condition. Significant differences are indicated as follows: n.s p≥0.05; *** p<0.001, Two-way ANOVA and post-hoc Tukey test. Scale bars: 50µm.

## Discussion

Here, we provide evidence that plants use the unique tension pattern at the organ surface of roots as input to regulate a cell-type specific growth control programme mediated by the small GTPase RAB-A5c. Through reinforcing cell walls at the organ surface^22^, RAB- A5c may allow epidermal cells to resist mechanical constraints, providing a molecular foundation for the previously observed role of the epidermis in constraining radial growth of the root^6,38^. Notably, our *in vivo* and *in silico* results suggest that cells in the lateral root cap can also mechanically constrain radial growth, purely through their geometry, which allows them to resist deformation even when exposed to high tensile stress. When root cap cover terminates within the meristematic region, maximum tension and thus radial growth control shifts to epidermal cells which activate RAB-A5c. While primary root meristems are fully covered by a root cap within 48h of germination, lateral root meristems remain partially uncovered for several days after their initiation, explaining the need for RAB-A5c-mediated growth control preferentially in lateral roots^21^.

Similar requirements for epidermis-specific reinforcement also exist in the outer-most cell layer of other plant organs^39,40^. RAB-A5c is expressed in the epidermis of shoots and leaf primordia and inhibition of its function causes defects in the morphogenesis of aerial organs, suggesting it may control epidermal growth in both roots and shoots^21^. We show here that RAB-A5c is expressed specifically in response to mechanical cues. Mechanical cues have been hypothesized to drive identity at the organ surface in shoots^23^, and recently, it has been shown that the activation of the core epidermal identity regulator AtML1 in mesophyll cells after wounding requires pressure release^35^, supporting the notion that pressure-driven tension can promote epidermal identity. Notably, a mechano-driven identity mechanism could contribute towards positional identity in plants, through which cells can re-establish cell layers with appropriate identities based on their tissue position rather than cell lineage^41,42^. In the root, epidermal cells of the same lineage experience differential mechanical tension depending on their position relative to the root cap. While some classic epidermal markers including *WEREWOLF* are expressed in both covered and uncovered epidermal cells^43^, others, including ARABIDOPSIS CRINKLY4^44^, are expressed exclusively in uncovered cells. This suggests that there are sub-populations of epidermal genes that follow different cues for expression, and raises the question whether mechanical cues control other epidermal functions, including barrier formation^45^ and sensing^46^.

## Material and Methods

### Plant materials and growth

The *Arabidopsis thaliana* ecotype Columbia=0 was used throughout. The following transgenic lines used in this study have been described before: *pRAB-A5c::YFP:RAB-A5c*^21^, *pRAB:A2a::YFP:RAB-A2a*^26^*, AtRPS5a>>Dex>>RAB- A5c[N125I]*^21^, *pUBQ10::LTi6B-TdTomato*^47^*, sombrero (smb)*^28^*, pUBQ10::YFP:NPSN12*^48^. All plants were grown at 20°C in a 16h:8h day:night cycle. Primary root age was counted from the moment of transfer to the growth chamber. Lateral roots were grown 8 - 12 days after germination on upright half-strength Murashige and Skoog medium (MS, Including Vitamins, Duchefa) plates with 2.5 mM MES hydrate (Sigma Aldrich), 1% w/v sucrose and 0.8% Difco^TM^ agar (BD Biosciences) at pH 5.7. For conditional expression using either dexamethasone or β-estradiol, seedlings were grown for 8 days from germination before transfer to half-strength MS medium containing either 1 µM Dex (Sigma Aldrich – diluted from a 10 mM stock in DMSO), 5 µM β-estradiol (Sigma Aldrich – diluted from 10 mM a stock in DMSO) or an equivalent volume of DMSO solvent for the indicated time period. Maximum root thickness was quantified as described before^49^.

### Molecular Cloning

The pSMB::XVE>>DT-A construct was generated using Gateway cloning (Invitrogen). The entry clone pEN-L4-pSMB-XVE-R1 contains a pSMB promoter fragment of 3071bp upstream of the translational start codon^24^ (upstream of the coding sequence of the chimeric transcription activator XVE^50^. The entry clone pEN-L1-DTA-L2 contains the coding sequence of the Diphtheria Toxin Fragment A (DT-A)^51^. These entry vectors were cloned into the destination vector pB7m24GW,3 using LR Clonase (Invitrogen) to create the expression vector pSMB::XVE>>DT-A.

To generate the pSMB:GRLhG4>>NAC46-BFP construct, the Golden Gate entry modules pGG-A-pSMB-B, pGG-B-GR-LhG4-E, pGG-E-35ST-F, and pGG-F-A-AarI-SacB- AarI-G-G were assembled in pFASTR-AG to create the destination vector pFASTR-pSMB- GR-LhG4-SacB. Entry modules pGG-A-pOP6-B, pGG-B-Linker-C, pGG-C-NAC046-D , pGG-D-P2A-mTagBFP2-NLS-E, pGG-E-G7T-F, and pGG-F-linkerII-G were inserted into this destination vector via a Golden Gate reaction resulting in the expression vector pFASTR-pSMB-GRLhG4>>pOP6:NAC046-P2A-mtagBFP2-NLS (short: pSMB:GRLhG4>>NAC46-BFP) . All vectors were collected from PSB plasmids stock (https://gatewayvectors.vib.be). Both expression vectors were transformed into Col-0 plants using the floral-dipping method^52^.

To generate the pUB::YFP-RABA5c line, the pUB promoter, YFP and RAB-A5c (At2g43130) sequences were respectively cloned into entry vectors pDONR™ P4-P1r, pDONR™ 221 and pDONR™ P2r-P3. The pUB promoter sequence contains the 1986 pb upstream of At4g05320 (UBQ10). The fluorescent variant sYFP2^53^ was used as YFP. For the cloning of the RABA5c entry vector, the primers attB2-RABA5c: GGGGACAGCTTTCTTGTACAAAGTGATGTCAGACGACGACGAGAGAGGCGAA and att3r-RABA5c: GGGGACAACTTTGTATAATAAAGTGAAAGAACTAATAATCACCACTACT were used on genomic DNA of *Arabidopsis thaliana* Col-0 with PrimeSTAR® Max DNA Polymerase (Takara Bio). For cloning, Gateway® BP Clonase™ II Enzyme Mix (Invitrogen) was used. The destination vector pUBQ10::sYFP2-RABA5c in pH7m34GW was obtained using the Gateway™ LR Clonase™ II enzyme mix (Invitrogen). Constructs were introduced into Agrobacterium tumefaciens strain C58 (pMP90) by electroporation, and Arabidopsis thaliana Col=0 plants were transformed by floral dip^52^.

### Confocal microscopy and image analysis

Confocal microscopy was performed using an inverted or upright Zeiss 980 CLSM using a C-Apochromat 40x/1.20 W Corr M27 objective. YFP, tdTomato, and PI were imaged as described before^26^. Image analysis and processing (orthogonal sectioning, maximum intensity projections, image assembly, surface renderings and quantification) was performed using Fiji^54^ or MorphoGraphX^55^.

### Laser ablation experiments

To perform laser ablation, 8 days old seedlings were placed in imaging chambers as detailed in^56^. Ablations were performed on lateral root tips using an inverted Leica DMI4000B spinning disc confocal microscope fitted with a pulsed UV laser from Teem Photonics (Grenoble, France), emitting at 355nm with a pulse duration of 500 ps, a pulse energy of 1 µJ, and a repetition frequency of 8 kHz, with a 40X/1.4NA C APO DICI objective and driven by MetaMorph and ILAS2 imaging software. Ablation area was adjusted to target 1-3 cells and 60% laser power with 200 repetitions was used. Confocal images were taken with an inverted Zeiss 980 CLSM using a C-Apochromat 40x/1.20 W Corr M27 objective before, immediately after and 6-24h after ablation. Between timepoints, plants were placed in the growth chamber. 3D segmentation and fluorescence quantification was performed as described below.

### 3D quantification of fluorescence

For 3D quantification of YFP:RAB-A5c or YFP:RAB- A2a fluorescence, CLSM stacks of ∼1mm long lateral roots containing the membrane channel (LTi6b-tdTomato) or PI were filtered using isotropic resampling and anisotropic filtering from the boundary_registration package (GitLab Project ID: 24938). Images were then analysed in MorphoGraphX using the CNN-UNet filter (BasselCombinedUNet.pt option), followed by ITK Auto-seeded 3D segmentation. After manual correction, 0.5µm meshes were built and smothered three times. The “Heat map classic” function was used to map volume and/or volumetric cell fluorescence from raw images and export data as csv files. Cell files were reconstructed manually by numbering cells based on their position relatively to the root cap end. Statistical analysis and plotting were performed using RStudio as described below.

### 2D quantification of fluorescence

For 2D quantification of *YFP:RAB-A5c* intensity, CLSM stacks of ∼1mm long lateral roots co-expressing *YFP:RAB-A5c* and *Lti6b-tdTomato* or PI were collected at Nyquist resolution (voxel size 99.5nm x 99.5nm x 550nm). Midplane longitudinal sections of meristematic cell files were generated in Fiji, and a line profile of 37 pixels thickness was plotted from the root tip through the centre of all epidermal cells in each section for *YFP:RAB-A5c* and the cell outline marker (*Lti6b- tdTomato* or PI). X positions and fluorescence intensities were imported as csv files into RStudio (https://www.rstudio.com/), cell boundaries were identified using an intensity- based threshold for the cell outline marker, and cell length and YFP fluorescence was quantified for each cell using a bespoke script.

### Volumetric shrinkage

To calculate the shrinkage of epidermal cell walls after turgor pressure release, 3D CLSM stacks of PI-stained lateral roots containing the YFP:NPSN12 plasma membrane marker were acquired first in water and then in 600mM Sorbitol. The difference in 3D volume was calculated by segmenting stacks in MorphoGraphX with the previously described method. To calculate crossectional shrinkage, cell dimensions in longitudinal, circumferential, and radial directions were determined using the 3DCellAtlas function in MorphographX^57^, circumferencial and radial lengths before and afterplasmolysis were multiplied to approximate crossectional area.

### Atomic Force Microscopy

10-12d old wild-type plants were immobilised to 60-mm petri dishes (Falcon 60 mm × 15 mm, Corning Ref. 351007) with the biocompatible glue Thin Pour (Reprorubber, Flexbar Ref 16135) and covered with MilliQ water after the glue was set (2-3 minutes). AFM experiments were performed on a stand-alone JPK Nanowizard III microscope (Bruker), driven by a JPK Nanowizard software 6.0. The acquisitions were performed with a spherical tip with a radius of 300nm (Biosphere NT_B300, Nanotools). The deflection sensitivity was calibrated by a force curve on a cleaned sapphire disk, while the spring constant provided by the manufacturer was directly used. The force curves were acquired using the following parameters: setpoint 1µN, Z length 2µm, speed 100µm/s. The acquisitions and were done using the Quantitative Imaging mode, with a typical image size of 30µmx50µm, with several tiles per root, followed by data analysis as described in^58^. Apparent Young’s moduli (Hertz model) were quantified on retract curves at different cell edges to reduce the effect of turgor on measurements, which is larger at cell faces. Young’s moduli maps were imported into FIJI^54^, cell edges were manually traced with a line thickness of 4 pixels, and average fluorescence was calculated for each edge.

### Serial Block-Face Scanning Electron Microscopy

For chemical fixation, roots from 10- 12d old wild-type seedlings were prepared following the protocol described in^59^: roots from 10-12-day old Arabidopsis plants were submerged in fixative solution containing 1% glutaraldehyde (v/v) and 0.5% paraformaldehyde (w/v) in 0.1 M sodium cacodylate buffer at pH 6.8 1 h at room temperature, post-fixed in 1% osmium tetroxide in 0.1 M sodium cacodylate buffer at pH 6.8 for 2h, and embedded in Spurr resin. Lateral roots were then mounted onto 3View stubs (Gatan) with conductive epoxy (Chemtronics) and were hardened for 4 h at 100 °C. The final trimmed block was sputter coated with gold for 30 s (layer thickness ∼20 nm) to improve conductivity. SBF-SEM images were collected on a Merlin Compact scanning electron microscope (Zeiss) with the Gatan 3View system. Section thickness was set to 100 nm, and the block face was imaged in variable pressure mode (∼50 Pa) at 4 kV acceleration voltage with a pixel dwell time of 7–8 μs and pixel size of 0.01 μm. Data processing (stack formation, image alignment, and trimming and scaling of sections to a common mean and SD) was done in the imod software package^60^. To quantify cell wall thickness, individual cross-section images were imported into Fiji, both sides of the cell wall for each cell face were traced manually, and XY Cartesian coordinates for each pixel on the outline trace were exported as csv files and imported into RStudio (https://www.rstudio.com/). For each pixel on one side, its closest neighbour on the other side was determined and the Euclidian distance between pixels calculated using the nn2 function in the RANN package (https://CRAN.R-project.org/package=RANN). Average distance between both pixels along the entire trace wall was calculated.

For high pressure freezing, lateral roots from 10-12-day old Arabidopsis plants were cut to <3 mm and placed in aluminium planchettes with 20% BSA in PBS as a cryoprotectant. High-Pressure Freezing was performed in a Leica ICE (Leica Microsystems) and transferred to an EM AFS2 freeze substitution system (Leica Microsystems) in cryovials containing 2% osmium and 0.1% uranyl acetate in acetone. Freeze substitution was performed over 90 hours, consisting of 12h at -90°C, a 6h transition period of -90°C - 85°C, a 48h transition period of - 85°C to -20°C, 12h at - 20°C and finally -20°C to 10°C over 12h. Samples were then washed in 100% acetone at room temperature and stained for an hour in 0.1% thiocarbohydrazide, before another wash in acetone and 1h incubation in 2% osmium in the dark. Samples were then washed in 100% acetone again, and infiltrated and embedded in epoxy resin (TAAB 812 Hard) of increasing concentrations (1:3, 1:1, 3:1, 1:0) in acetone over 4 days, before polymerization in 100% resin at 60°C for 48h. Resin blocks containing root tips were trimmed and mounted onto aluminium pins using conductive epoxy glue and Silver DAG 1415M. Blocks were sputter coated with a 10-13 nm layer of gold using an Agar Auto Sputter Coater (Agar). A Merlin VP compact high-resolution scanning electron microscope (Zeiss) equipped with a 3View 2XP stage, an OnPoint back-scattered electron detector (Gatan-Ametek) and a focal charge compensation device (Zeiss) were used for SBF-SEM image generation. For image capture, the following setting were used; 1.8 kV accelerating voltage, 30 µm aperture, high vacuum, 100% focal charge compensation, 5 nm pixel size, 50 nm section thickness, 1 µs pixel time and approximately 12000 x 13000-pixel image size.

### Computational models

We considered the 2D model of pressurized cells with cell walls made of a linear elastic material on a domain Ω. The idealized cellular geometry of the domain Ω corresponding to the circular root cross section (Figure S8A) was defined parametrically, with parameter values (number of layers and cells per layer, layer heights, internal and external cell wall thickness, curvature radii in cell corners) inspired from measurements (Figure 3). Each cell in the tissue is considered as a homogeneous subdomain, and the material properties (Young’s modulus for instance) might vary from cell to cell. Geometric and mechanical parameter values are given in the legend of Figure S8A. This domain is considered as the stress-free reference configuration of the system, that can deform with a deformation field u and strain tensorfield ε. The stress tensor σ is related to the strain by the linear constitutive law σ =H ε , with H the four rank elasticity tensor field, that is isotropic and cellularily piecewise constant, encoding as parameters a uniform Poisson ratio and a Young’s modulus that can vary from cell to cell. A cellularily piecewise constant pressure field P is also considered on the domain, that encodes in particular the pressure values at the interior boundary Γ of each cell (FigureS8A). Then, the mechanical equilibrium after pressurizing the cells is given by the following PDE for the displacement u with n the unit normal on the boundary.

We used finite element method and in particular the open source software FreeFem++ [*] to solve this equation, find the deformed domain (Figure S8B) and compute tensile and compressive stress maps in the cell walls (Figure S8B) when cells are pressurized. Tensile stress values per cell (Figures 3G, 3M, 4A-B, 5B, 5D, S4 A-B) are computed by taking the mean of these scalar fields per cell domain. The simulation script is accessible at the repository https://gitlab.inria.fr/mosaic/publications/layers.

Model parameters:

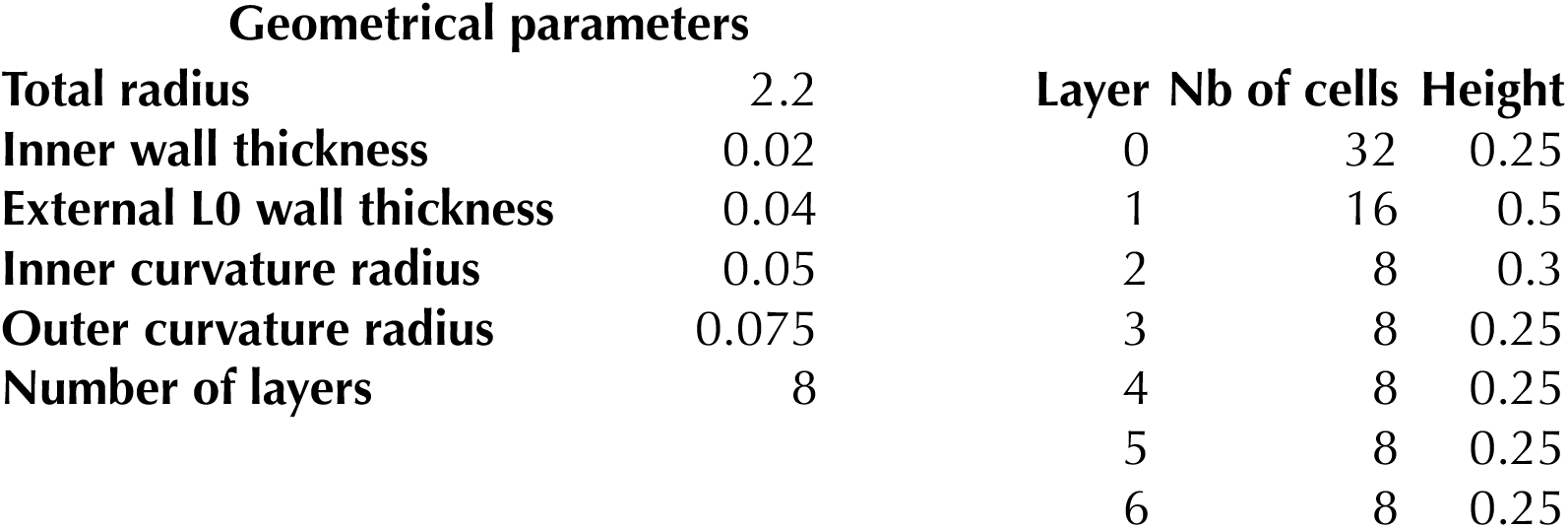
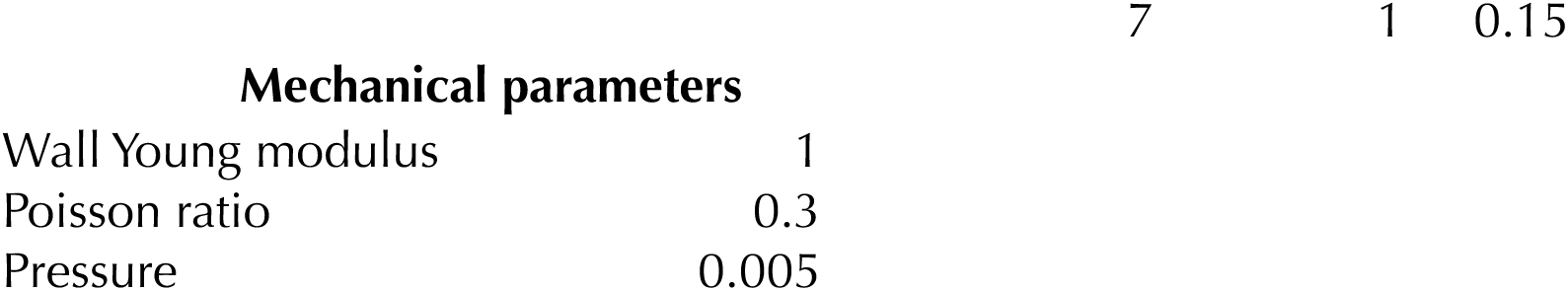

### Statistical data analysis and plotting

The stats package in R was used for statistical analysis^54^. Two-way ANOVA (analysis of variance) was performed using the aov function, Tukey’s test was performed using the TukeyHSD function and Student’s t-test were performed using the t.test function. Box- and Ribbon-plots were generated in R using the ggplot2 function^61^. In box plots, the median is displayed as a line, lower and upper hinges correspond to the 25th and 75th percentiles, the lower and upper whiskers extend from the hinge to the smallest or largest value no further than 1.5 * IQR from the hinge. Data points were also plotted individually. Ribbon plots show the data mean +/- standard deviation (shaded areas).

All experiments were conducted at least twice independently. For experiments involving confocal images of lateral roots, 3-8 lateral roots were imaged for each condition/genotype in each experimental repeat, for experiments involving bright-field images, 18 – 30 lateral roots were imaged from each condition/genotype. Data from experimental repeats were pooled where specified, else data from one representative experiment are shown.

## Acknowledgments

We acknowledge funding from the Leverhulme Trust (Early Career Fellowship ECF-2017- 483 to CK), the European Research Council (ERC-2020-Stg 948514 - EDGE-CAM to CK), and the ANR France 2030 administered by the INRAE EXPLOR’AE programme (MechanoID to CK). We acknowledge the contribution of SFR Biosciences (Universite Claude Bernard Lyon 1, CNRS UAR3444, Inserm US8, ENS de Lyon) PLATIM-LyMIC, especially Elodie Chatre for assistance on the ablation experiment. We thank Yvon Jallais for sYFP2 and pUBQ10 entry vectors. We warmly thank Louise Hughes and Chris Hawes for help with SBF-SEM experiments. We thank Benoit Landrein for many stimulating discussions during the genesis of this project.

## Author contributions

Conceptualization: ZNV, CK

Methodology: ZNV, AK, SB, MM, CL, TV, QF, FdW, RH, MN, CK

Investigation: ZNV, AK, SB, NG, NF Visualization: ZNV, AK, SB, CK Funding acquisition: CK

Project administration: CK Supervision: VK, MN, CK Writing – original draft: CK

Writing – review & editing: ZNV, CK

## Competing interests

Authors declare that they have no competing interests.

Materials and correspondence: Please contact Charlotte Kirchhelle for the materials used in this study.

## Notes

### Competing Interest Statement

The authors have declared no competing interest.

