## supplementary figures for "Plant cells at the organ surface use mechanical cues to activate a specific growth control programme"

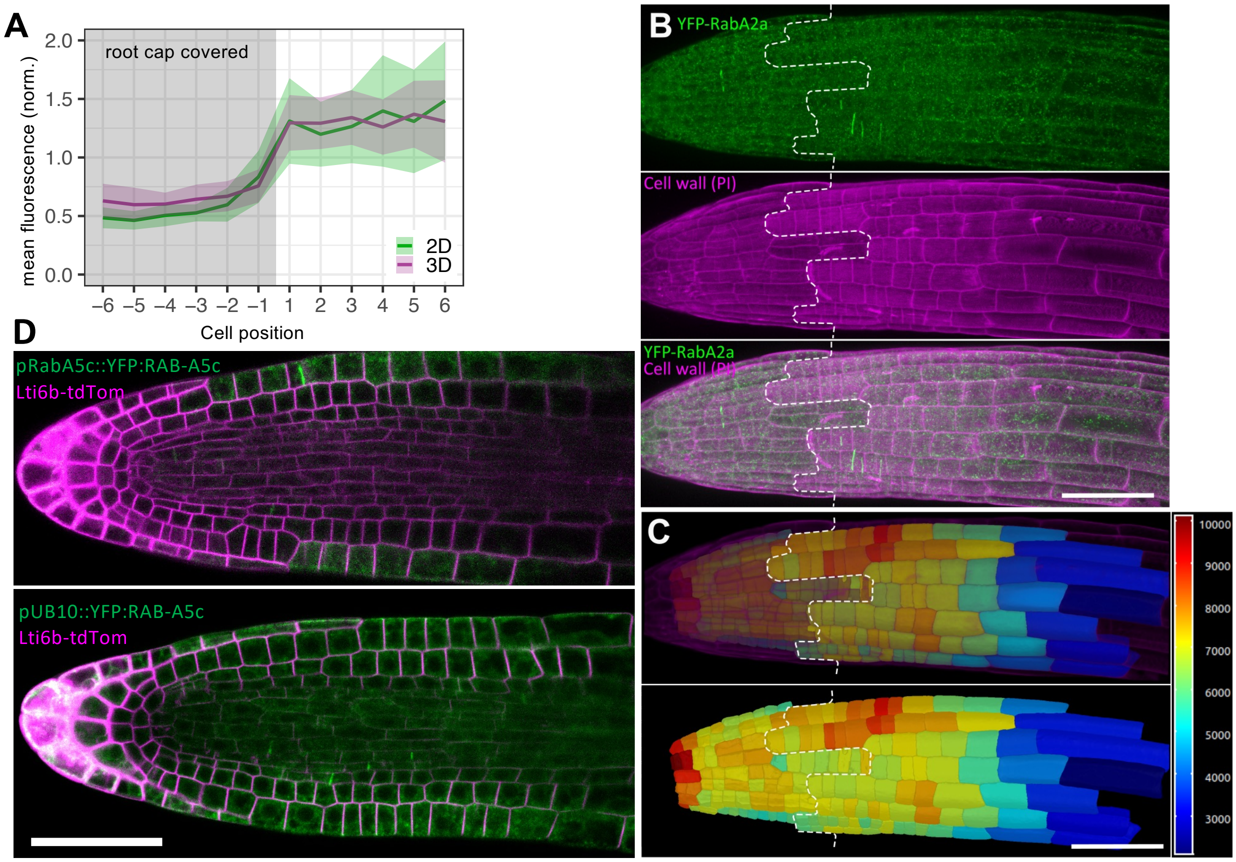


**Figure S1: Expression patterns of Rab-A GTPases in lateral roots. (A)** Mean fluorescence intensity of YFP:RAB-A5c per cell along epidermal cell files of lateral roots quantified in 2D on midplane sections or in 3D through cell segmentation. Fluorescence was quantified along 3-5 individual cell files per root, cell files were aligned based on the position of the root cap, with the last covered cell labelled as -1 and the first uncovered cell labelled as 1. Ribbon plots represent average fluorescence +/- 1SD for 14 cell files from 3 roots. **(B)** Maximum intensity projection of a confocal laser scanning microscopy (CLSM) stack of a lateral root expressing YFP:RAB-A2a stained with the cell wall marker propidium iodide (PI). Dashed line indicates the end of the root cap. **(C)** MorphoGraphX 3D segmentation of epidermal cells in the lateral root shown in (B). Heat map depicts mean fluorescence intensity of YFP:RAB-A2a. Dashed line indicates the end of the root cap. **(D)** XY section of a CLSM stack of lateral roots expressing the membrane marker Lti6b-TdTomato and YFP:RAB-A5c under either its own promoter (*pRAB-A5c::YFP:RAB-A5c*) or the ubiquitin10 promoter (*pUB10::YFP:RAB-A5c*). Scale bars: 50 µm.

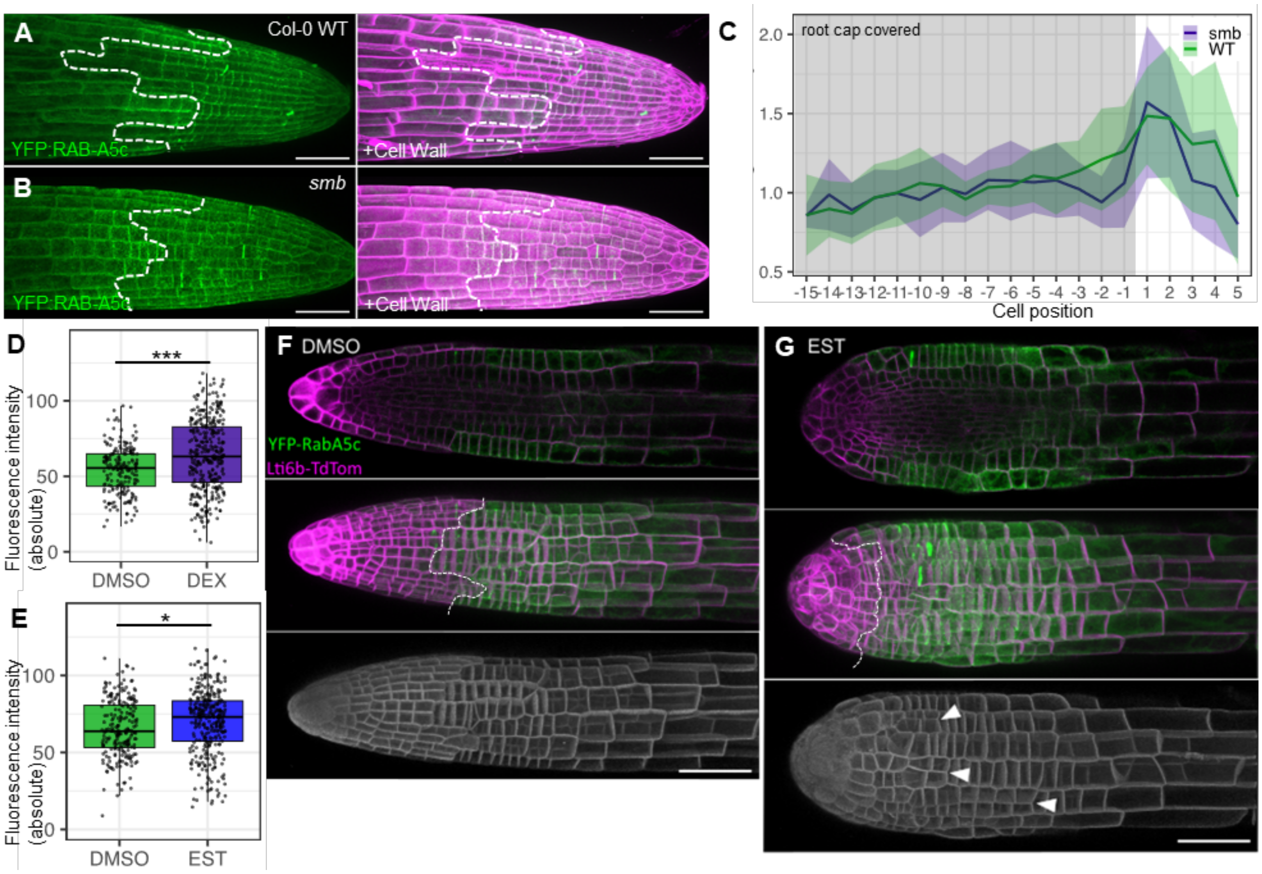


**Figure S2: YFP:RAB-A5c is expressed in cells at the organ surface. (A,B)** Maximum intensity projections of CLSM stacks of 36h old primary roots expressing YFP:RAB-A5c stained with the cell wall marker PI in a wild type (A) and *sombrero* (B) background. Dashed line indicates the end of the root cap. **(C)** Mean fluorescence intensity of YFP:RAB-A5c per cell along epidermal cell files of primary roots such as those shown in (A,B). Fluorescence was quantified in 2D, cell files were aligned based on the position of the root cap, with the last covered cell labelled as -1 and the first uncovered cell labelled as 1. Ribbon plots represent average fluorescence +/- 1SD. N=10 (WT) and 12 (*smb*) cell files from 3 roots. **(D)** Box plot of average YFP:RAB-A5c intensity in uncovered cells of *pSMB:GRLhG4>>NAC46-BFP* lateral roots after 48h on DMSO (n=200) or DEX (n=346). Student's T test; *** $p<0.001$ **(E)** Box plots of average YFP:RAB-A5c intensity in uncovered cells of *pSMB:XVE>>DT-A* #1 lateral roots after 48h on DMSO (n=225) or EST (n=311). Student's T-test; * $p<0.05$. **(F,G)** XY midplane optical sections (top), maximum intensity projections (middle) and MorphoGraphX renderings (bottom) of CLSM stacks from lateral roots co-expressing YFP:RAB-A5c, Lti6b:tdTomato, and an independent transgene insertion of Estradiol (Est)-inducible *pSMB:XVE>>DT-A* #2 after 48h treatment with DMSO as a control (F) or 5µM Est to induce lateral root cap death (G). Arrowheads indicate misplaced longitudinal divisions. Dashed line indicates the end of the root cap. Scale bars: 50µm

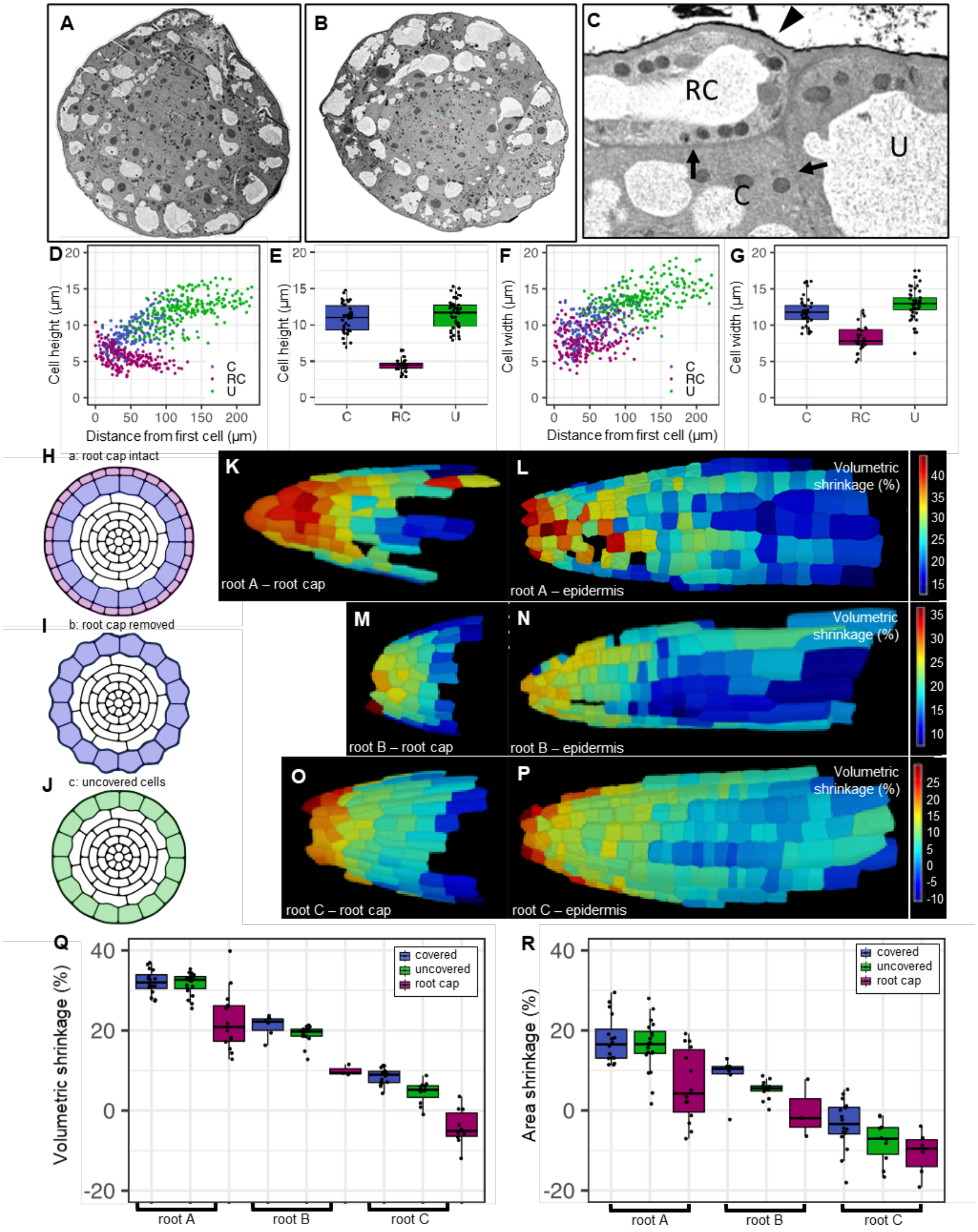


**Figure S3: Geometric and mechanical characterisation of lateral root cells. (A-C)** Transverse cross-sections (A,B) and a magnified view (C) of a serial block-face scanning electron micrograph (SBF-SEM) of a cryo-fixed wild-type lateral root. Note the outer epidermal face (arrowhead) is thicker than internal cell faces (arrows) as in the PFA-fixed root (Figure 3A-C), indicating relative cell wall thicknesses are well-preserved though PFA fixation. **(D-G)** Cell radial lengths (Cell height) (D,E) and circumferential lengths (Cell width) (F,G) from three 3D-segmented wild-type lateral roots. (E,G) are box plots of cells from each root in (D,F) that were in the zone between the first uncovered and last covered cell. **(H-J)** Deformed configurations of computational models of lateral root cross-sections corresponding to cases a,b, and c presented in Figure 3F. **(K-P)** Volumetric shrinkage in the root caps (K,M,O) and epidermis (L,N,P) of three roots after osmotic treatment with 600mM sorbitol to release turgor pressure. **(Q,R)** Volumetric shrinkage (Q) and Area shrinkage (R) of cells in roots shown in (K-P).

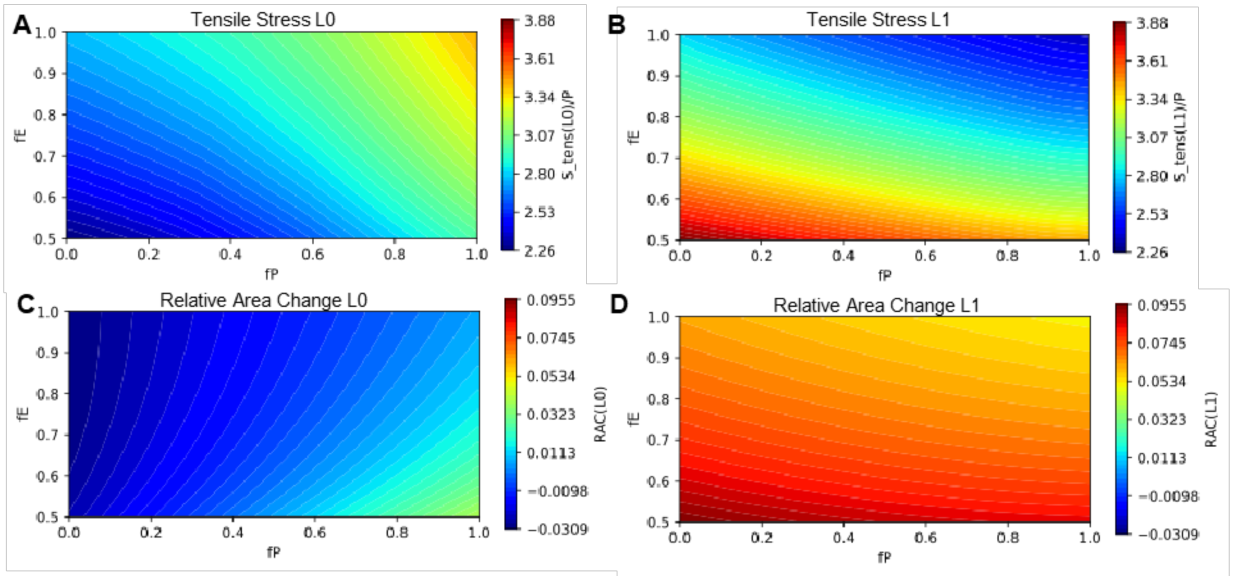


**Figure S4: Stress and Strain in L0 and L1 layers of root cross-section models.** Maps of tensile stress (A,B) and relative cross-sectional area change (C,D) in the root cap (L0; A,C) and covered epidermis (L1; B,D) in response to changes in turgor pressure P and elastic modulus E within the L0 layer.

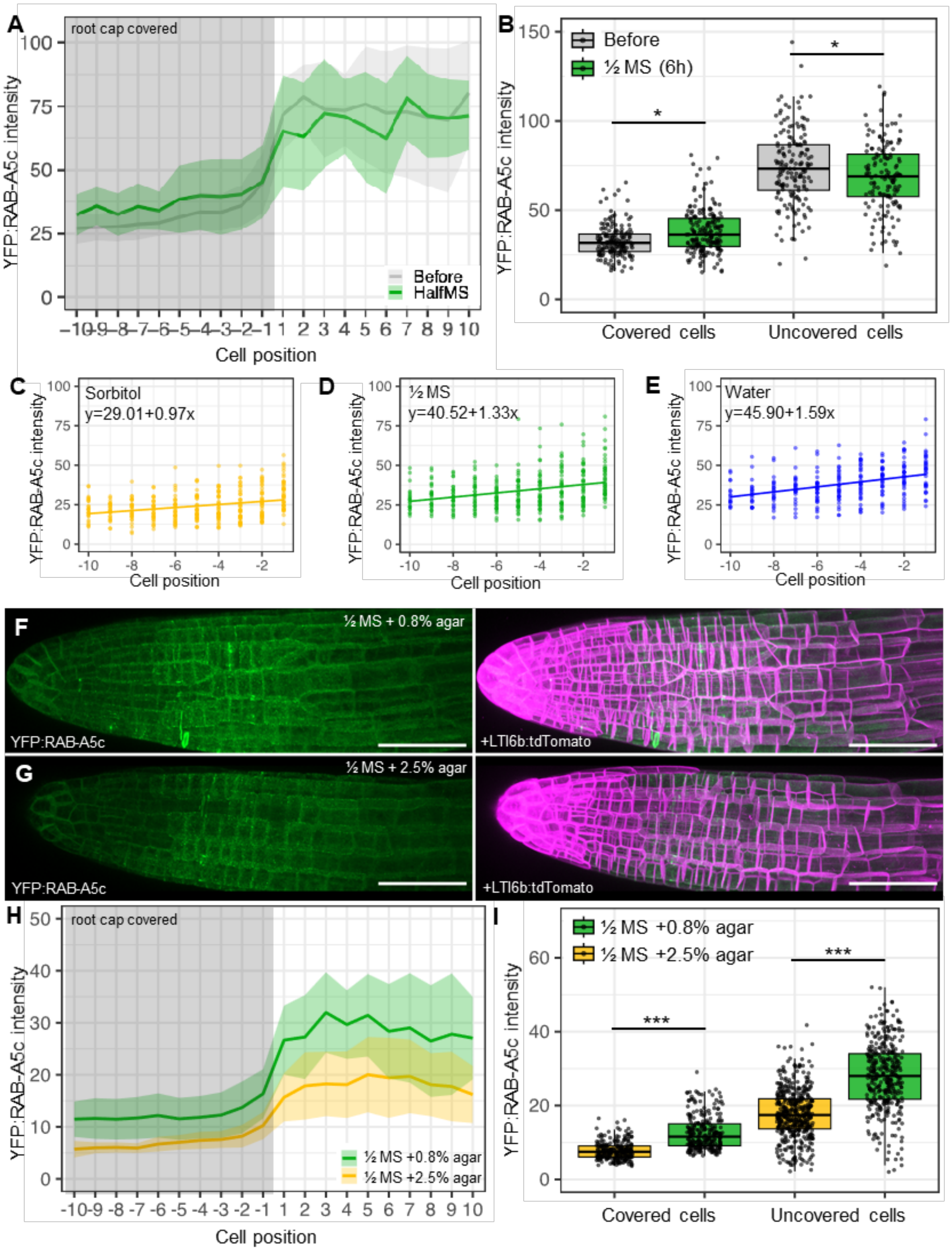


**Figure S5: YFP:RAB-A5c levels can be modulated through changing tissue tension. (A)** Mean fluorescence intensity of YFP:RAB-A5c per cell along epidermal cell files of lateral roots before or after 6h submersion in liquid ½ MS medium. N=20 (Before), 22 (HalfMS) cell files from 4 roots. Fluorescence was quantified in 2D, cell files were aligned based on the position of the root cap, with the last covered cell labelled as -1 and the first uncovered cell labelled as 1. Ribbon plots represent average fluorescence +/- 1SD. **(B)** Box-plot showing YFP:RAB-A5c intensity in covered and uncovered cells before or after 6h submersion in liquid ½ MS medium. N≥149. **(C-E)** Linear model (LM) fit of fluorescence in covered epidermal cells from roots such as those shown in Figure 4C-E. Note gradient is lowered in the presence of sorbitol and increased in the presence of water compared to ½MS controls. **(F,G)** Maximum intensity projection of CLSM stacks of lateral roots expressing YFP:RAB-A5c and Lti6b:tdTomato grown on ½ MS + 0.8% agar (F) or ½ MS + 2.5% agar (G). **(H)** Mean fluorescence intensity of *YFP:RAB-A5c* per cell along epidermal cell files of lateral roots such as those shown in (F,G). N=42 (0.8% agar), 56 (2.5% agar) cell files from 11 and 14 roots respectively. Fluorescence was quantified in 2D, cell files were aligned based on the position of the root cap, with the last covered cell labelled as -1 and the first uncovered cell labelled as 1. Ribbon plots represent average fluorescence +/- 1SD. **(I)** Box plots of average YFP:RAB-A5c intensity in covered and uncovered cells from data shown in (H). N= 238 (0.8% agar covered), 247 (2.5% agar covered), 338 (0.8% agar uncovered), 444 (2.5% agar uncovered). Significant differences in thickness are indicated as follows: * $p<0.05$; *** $p<0.001$, Two-way ANOVA and post-hoc Tukey test. Scale bars: 50µm

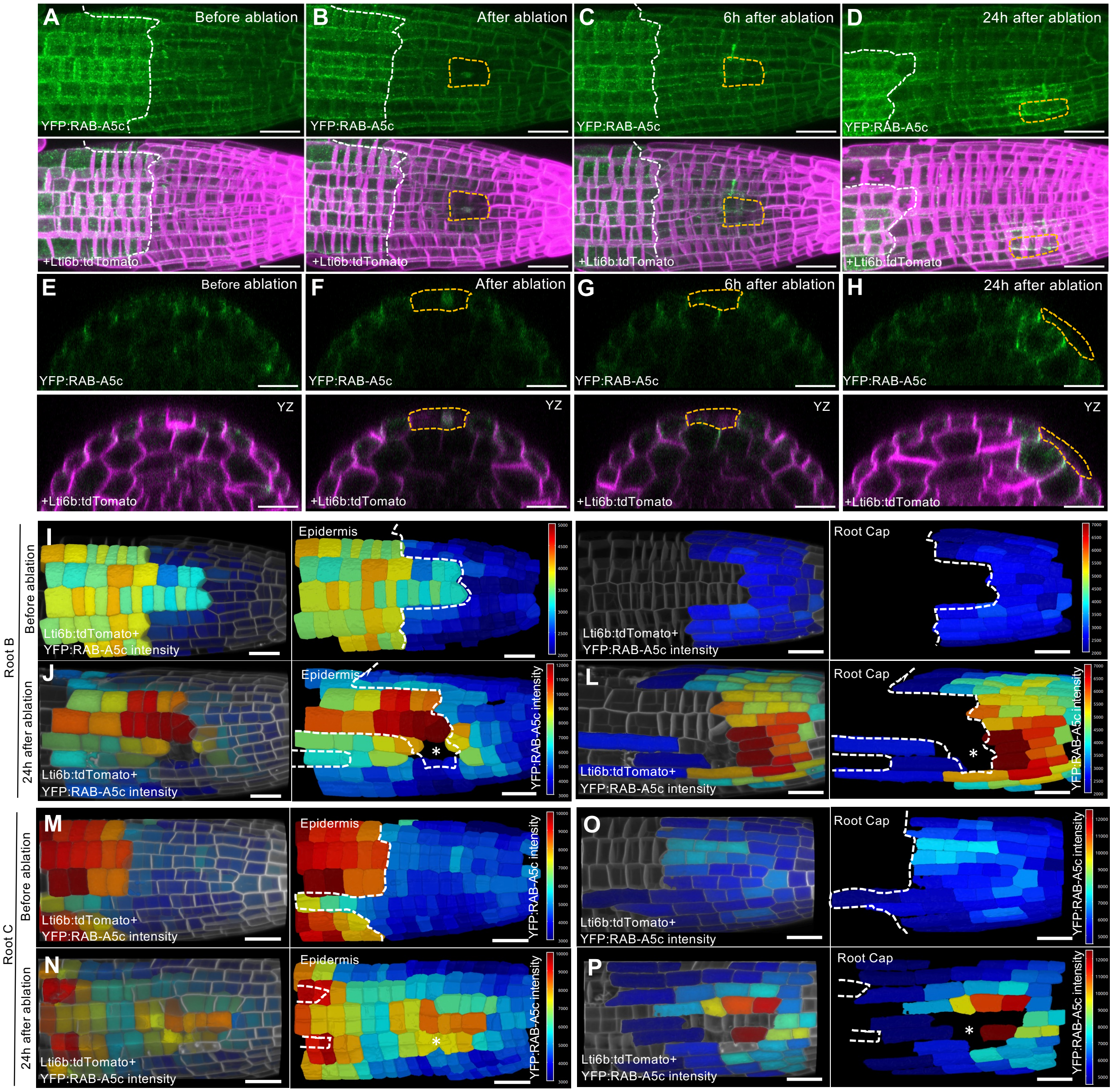


**Figure S6: YFP:RAB-A5c levels can be locally modulated through cell ablations. (A-H)** Maximum intensity projections of CLSM stacks (A-D) and YZ sections (E-H) of a lateral root co-expressing YFP:RAB-A5c and Lti6b:tdTomato before and immediately after, 6h, and 24h after ablation of a lateral root cap cell. Dashed white line indicates end of the root cap, dashed yellow line indicates the ablation site. Note pronounced photobleaching of both fluorophores beneath the ablation site immediately after and 6h after ablation. **(I-P)** MorphoGraphX volumetric fluorescence intensity maps of meristematic cells (I,J,M,N) and root cap cells (K,L,O,P) from two lateral roots co-expressing YFP:RAB-A5c and Lti6b:tdTomato before and 24h after ablation in the root cap and epidermal layers. Note expression is activated in epidermal and root cap cells surrounding the wound site. Scale bars: 20µm.

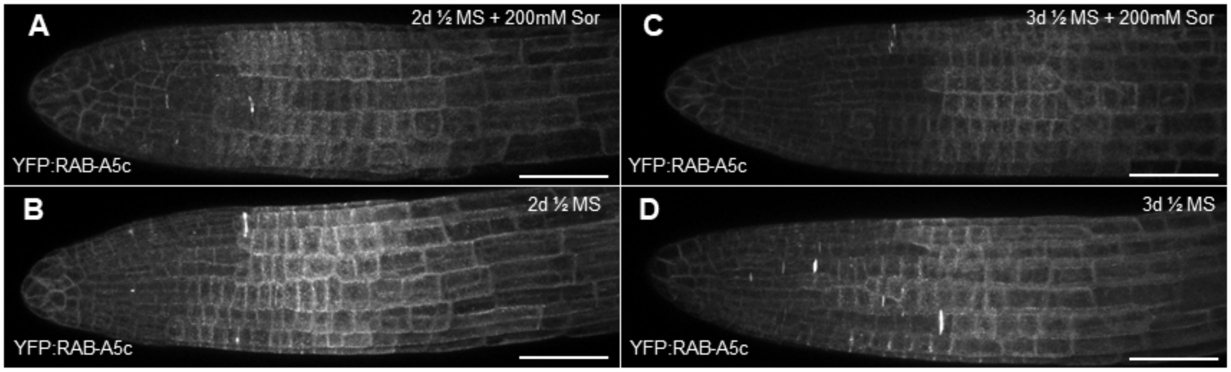


**Figure S7: YFP:RAB-A5c levels in response to long-term treatments with Sorbitol.** Maximum intensity projections of CLSM stacks of lateral roots expressing YFP:RAB-A5c after 2d (A,B) and 3d (C,D) on ½ MS (B,D) or ½MS supplemented with 200 mM sorbitol (A,C). Scale bars 50µm.

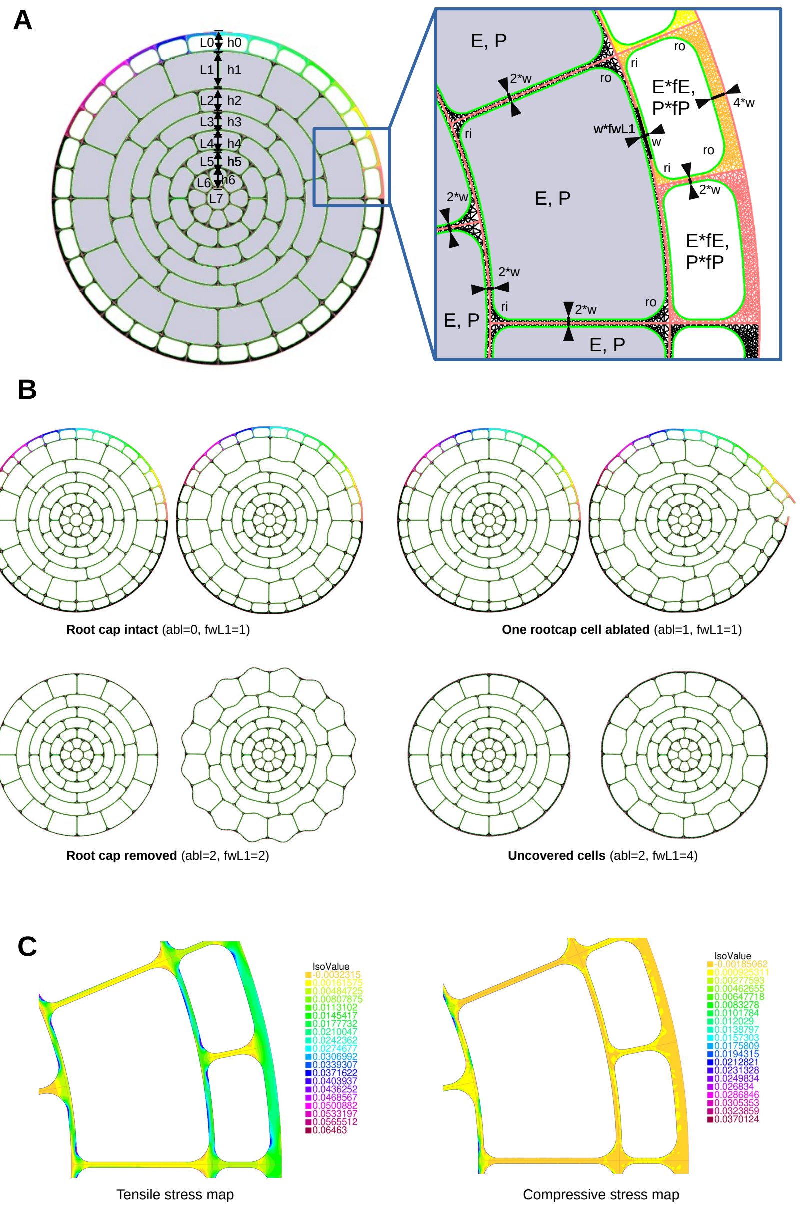


**Figure S8: Computational models. (A)** Geometrical and mechanical parameters. The circular root cross section of radius 2.2 is constructed from 8 ring cellayers (of heights 0.25, 0.5, 0.3, 0.25, 0.25, 0.25 and 0.15) as is shown. The half inner wall thickness is w=0.01. The external cell wall thickness of L0 is 4*w. Curvature radii in the inner corners are ri=0.05 while in the outer corners are ro=0.075. The outer cellwall thickness of the L1 cells can be tuned by a factor fwL1. Each cell is a subdomain in the domain of the whole tissue, and a cellularily piecewise constant Young's modulus and pressure field is defined. Here we consider uniform Young's modulus E=1 and P=0.005 in all interior layers (gray cells), while Young's moduli and/or pressure in cells of the outer L0 layer (white cells) can be tuned by a factor fE and fP with respect to these inner values. **(B)** Geometry of the four situations considered in this manuscript: „Root cap intact" contains all the layers and the wall between the L0 and L1 layers is of thickness 2w, as all other interior walls. For the „One rootcap cell ablated" situation the outer cell wall of one cell is cut and the pressure in this cell is 0. The tissue corresponding to „Rootcap removed" and „Uncovered cells" has all but the outer L0 layer, they differ only in the thickness of the outer L1 wall thickness, tuned by the fwL1 parameter. The parameter „abl" in the simulation script allows to use the above presented three geometries. **(C)** Tensile and compressive stress maps. Using the local principal stress values σ1 and σ2, tensile and compressive stresses are defined as S_tensile=(σ1)++(σ2)+ and S_compressive=(-σ1)++(-σ2)+ with ()+ denoting the positive part function.
